# Paternal morphine exposure enhances morphine self-administration and induces region-specific neural adaptations in reward-related brain regions of male offspring

**DOI:** 10.1101/2023.01.03.522600

**Authors:** Andre B. Toussaint, Alexandra S. Ellis, Angela R. Bongiovanni, Drew R. Peterson, Charlotte C. Bavley, Reza Karbalaei, Hannah L. Mayberry, Shivam Bhakta, Carmen C. Dressler, Caesar G. Imperio, John J. Maurer, Heath D. Schmidt, Chongguang Chen, Kathryn Bland, Lee-Yuan Liu-Chen, Mathieu E. Wimmer

**Affiliations:** Department of Psychology, Program in Neuroscience Temple University, Philadelphia, PA, USA; Department of Psychiatry and Behavioral Science, Temple University, Philadelphia, PA, USA; Department of Biobehavioral Health Sciences, School of Nursing, University of Pennsylvania, Philadelphia, PA, USA; Department of Psychiatry, Perelman School of Medicine, University of Pennsylvania, Philadelphia, PA, USA; Center for Substance Abuse Research and Department of Neural Sciences. Temple University Lewis Katz School of Medicine, Temple University, Philadelphia, PA, USA

**Keywords:** Epigenetic inheritance, multigenerational, mu-opioid receptor, RNA sequencing, behavior, addiction

## Abstract

**Background:** A growing body of preclinical studies report that preconceptional experiences can have a profound and long-lasting impact on adult offspring behavior and physiology. However, less is known about paternal drug exposure and its effects on reward sensitivity in the next generation.

**Methods:** Adult male rats self-administered morphine for 65 days; controls received saline. Sires were bred to drug-naïve dams to produce first-generation (F1) offspring. Morphine, cocaine, and nicotine self-administration were measured in adult F1 progeny. Molecular correlates of addiction-like behaviors were measured in reward-related brain regions of drug naïve F1 offspring.

**Results:** Male, but not female offspring produced by morphine-exposed sires exhibited dose-dependent increased morphine self-administration and increased motivation to earn morphine infusions under a progressive ratio schedule of reinforcement. This phenotype was drug-specific as self-administration of cocaine, nicotine, and sucrose were not altered by paternal morphine history. The male offspring of morphine-exposed sires also had increased expression of mu-opioid receptors in the ventral tegmental area but not in the nucleus accumbens.

**Conclusions:** Paternal morphine exposure increased morphine addiction-like behavioral vulnerability in male but not female progeny. This phenotype is likely driven by long-lasting neural adaptations within the reward neural brain pathways.

## Introduction

The opioid crisis remains a massive public health concern in the United States. Since the beginning of the 21^st^ century, increases in dispensing rates of opioids such as morphine have resulted in a rapid growth in diagnosed opioid use disorder [1, 2]. As rates of opioid misuse and abuse increase, so has the attention towards understanding the long-term effects on subsequent generations. Parental insults to stress [3, 4], diet [5, 6], and environmental toxins [7] have been extensively studied, revealing significant long-lasting impacts on future generations. There is also a growing body of literature showing that parental exposure to drugs of abuse may influence drug abuse liability in their offspring [8–18].

For prenatal opioid exposure, many of the aforementioned studies have focused on the maternal lineage. Maternal exposure to morphine has been shown to both increase [19] and decrease [12] risk towards abuse of opioids in offspring. Prenatal opioid exposure can also impact memory, anxiety, and pain sensitivity in offspring [19–26]. The route of administration for opioids is an important determinant in abuse liability of drugs [27, 28] and is therefore an important consideration when parsing this literature. To date, only one study has examined the impact of paternal morphine exposure on self-administration of opioids in offspring [29]. Other studies have investigated sensitivity to opioids in morphine-sired offspring using the two-bottle choice paradigm [30] or conditioned placed preference [31]. Here, we use a highly translational rodent model of paternal morphine self-administration to investigate vulnerability to drug taking in offspring.

In summary, we examined the effects of paternal morphine self-administration on addiction-like behaviors using self-administration of morphine, cocaine, and nicotine in offspring. Molecular correlates were measured using mu-opioid receptor binding in drug-naïve saline-sired and morphine-sired offspring.

## Methods and Materials

### Subjects

Male Sprague Dawley rats (250 – 300g) from Taconic Biosciences (Hudson, NY, USA) were procured and paired-housed before and after intravenous catheterization surgery for the parental (F0) generation. First generation progeny (F1) were bred in house (details below). All rats were kept in a temperature and humidity-controlled room on a 12h light/dark cycle, with lights off at 8:30 am. Food and water were available *ad libitum*. All procedures in the F0 and F1 generations were carried out in accordance with the Institutional Animal Care and Use Committee (IACUC) guidelines of Temple University.

### Drugs

Morphine sulfate, cocaine hydrochloride, and nicotine hydrogen tartrate were obtained from Spectrum Chemical (Gardena CA) and dissolved in sterile 0.9% saline.

### Behavioral Procedures

#### Catheterization Surgery

Adult rats were anesthetized using an i.p. injection of a ketamine/xylazine cocktail (80 and 12 mg kg ^-1^, respectively). An indwelling silastic catheter (Instech Laboratories, Inc. Plymouth Meeting, PA) was implanted into the right jugular vein, sutured in place, and mounted dorsal to the shoulder blade using a back-mount platform. Catheters were flushed daily with timentin (0.93 mg ml^-1^), dissolved in heparinized saline and sealed using metal caps, when not in use. Following catheterization surgery, animals were recovered for 1 week prior to the start of self-administration.

#### F0 morphine self-administration

Sires were allowed to lever press for intravenous morphine (daily 3-hr sessions, 0.75 mg/kg morphine/59 μl saline, infused over 5 s) for 60 consecutive days (duration of rodent spermatogenesis) on a fixed ratio 1 (FR1) schedule of reinforcement, where one lever press resulted in a single infusion of morphine. Age-matched control rats received saline over the same time period. We and others have previously used this model [32, 33]. Twenty-four hours after the last of the 60-day morphine self-administration session, sires were pair-housed with drug-naïve dams for up to 5 days to copulate. Both groups of sires continued to self-administer morphine (or saline) during the mating period to avoid withdrawal-mediated anxiety and stress as a confounding factor [34–36]. At PND 21, we weaned offspring derived from morphine-experienced sires (morphine-sired) and saline controls (saline-sired) and pair-housed them with same-sex littermates. For each of the four groups (male saline-sired, male morphine-sired, female saline-sired, female morphine-sired) 1-2 adult (post-natal day 60 - 120) rats were randomly selected for behavioral and molecular analyses (**Figure 1A**).

**Figure 1.**
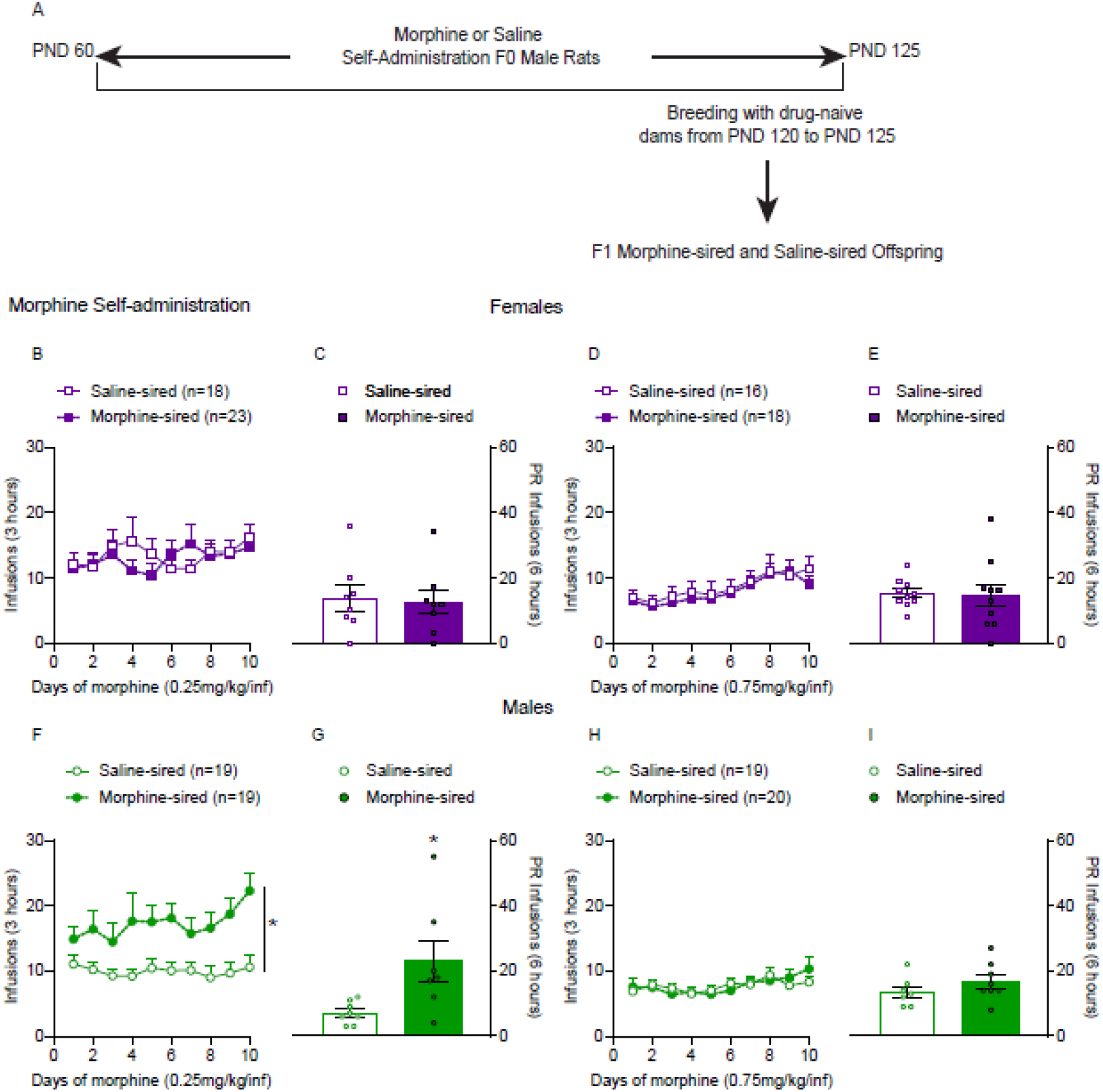
Increased self-administration and reinforcing efficacy of low dose morphine in morphine-sired adult male rats. **A.** F0 males self-administered either morphine (0.75 mg/kg/infusion) or saline for 60 days consecutively. On day 61, sires were paired with drug-naïve dams for breeding for up to 5 days. Following breeding, dams were separated from sires, and F1 progeny were born. **B.** Paternal morphine consumption did not alter the acquisition of morphine self-administration in adult female rats (from 8 saline-exposed sires and 10 morphine-exposed sires) at the 0.25mg/kg dose. **C.** Under the progressive schedule of reinforcement, saline-sired (from 8 sires) and morphine-sired (from 10 sires) female rats took similar amounts of morphine infusions. **D.** At the 0.75mg/kg dose, morphine self-administration was intact in female rats (from 8 saline-exposed sires and 6 morphine-exposed sires). **E. T**here was no difference between saline-sired (from 8 sires) and morphine-sired (from 6 sires) female rats when they were switched to a progressive ratio schedule. **F.** In contrast to female progeny, paternal morphine exposure significantly enhanced morphine self-administration at the low dose in morphine-sired adult male rats (from 8 saline-exposed sires and 8 morphine-exposed sired). **G.** In line with these results, morphine-sired (from 8 sires) rats took significantly more infusions of morphine compared to saline-sired rats (from 8 sires) under a progressive ratio schedule. **H.** There was no difference in morphine infusions between saline-sired (from 5 sires) and morphine-sired male rats (from 6 sires) at the 0.75 mg/kg dose. **I.** Paternal morphine consumption did not alter morphine self-administration at the higher dose in male rats (from 5 saline-exposed sires and 6 morphine-exposed sires) under a progressive ratio schedule of reinforcement. Data are expressed as the mean ± S.E.M. *p < 0.05.

#### Morphine self-administration in F1 offspring

Naïve male and female F1 offspring were allowed to self-administer morphine at two different doses: 0.25 or 0.75 mg/kg/infusion for 10 days on a FR1 schedule of reinforcement during daily 3-hour sessions. On day 11, rats were switched to a progressive ratio (PR) schedule in which the response requirement for each subsequent morphine infusion increased over the course of a 6-hr hour time period. We also collected and analyzed vaginal cytology samples from female progeny to determine whether the estrous cycle may influence morphine self-administration (for details, see supplemental methods). For all morphine self-administration experiments, we used a progressive ratio responding schedule that has been shown to support morphine self-administration consistently over consecutive days [37, 38]. The number of lever pressing responses required to receive an infusion increased over the course of the session using the following equation: Response requirement = Round (C1 x e^[C2 x (Step Number C3)]^ – C1 + C4) where results are rounded to the nearest integer value; step number is the number of ratios completed and C1, C2, C3 and C4 are constants with values of 10, 0.035, 1, and 0.5, respectively. The response requirement increased according to this sequence: 1, 1, 1, 2, 2, 2, 3, 3, 3, 4, 4, 5, 5, 6, 6, 7, 7, 8, 9, 9, 10, 11, 11, 12, 13, 14 etc.

#### Cocaine Self-Administration in F1 offspring

A separate drug-naïve group of littermates derived from morphine- and saline-exposed sires were used in this experiment. Adult F1 rats self-administered cocaine at two different doses (daily 2hr session, 0.25 or 0.5 mg/kg) on an FR1 schedule of reinforcement for 10 consecutive days. Each cocaine infusion was followed by a 20-s timeout period, during which cocaine was not infused but responses were recorded. Following 10 days on FR1, rats were switched to a progressive ratio schedule of reinforcement where the response requirement - (R(i)= [5e^0.2i^–5]) - expired when a rat took more than 30 minutes to lever press for the subsequent cocaine infusion.

#### Determination of mu-opioid receptor (MOR) expression and MOR-mediated G protein activation in reward-related brain regions of drug-naïve offspring

MOR expression was measured with binding of the selective agonist [^3^H]DAMGO and MOR-mediated G protein activation was determined with DAMGO-stimulated [^35^S]GTPγS binding per our published procedures [39] with some modifications.

NaïveAdult (60+ days) naïve first-generation saline- and morphine-sired offspring were sacrificed by decapitation and brains removed and frozen immediately. Bilateral VTA and NAc punches were collected from frozen brains. NAC from one rat or VTAs from two rats were pooled as one sample, put into an Eppendorf tube and kept frozen until experiments.

Frozen rat VTA tissues were thawed on ice, sonicated to disrupt cells in a total volume of 100 μl [^3^H]DAMGO binding buffer (50 mM Tris buffer containing 0.2mM phenylmethylsulfonyl fluoride, pH7.4) per tube immediately before the binding assays. Because of the small size of the samples, we did not prepare membranes. Similary, two NAc tissue punches (1 mm) from one rat were placed in 1.5ml Eppendorf tubes and immersed in ice-cold 200uL binding buffer per tube. The tissues were homogenized by sonication for five short bursts. An aliquot of 10uL was taken from each sample for determination of protein contents by bicinchoninic acid assay (BCA assay) followed by of Nanodrop measurement.

MOR radioligand binding was performed with 1.5n M [^3^H]DAMGO in 50 mM Tris buffer containing 0.2mM phenylmethylsulfonyl fluoride, pH7.4 in a final volume of 1 ml and naloxone (10 μM) was used to define non-specific binding. Binding was done in duplicate for the NAc samples, but in singleton for the VTA samples because the VTAs were very small. Binding interaction was allowed to reach equilibrium by incubation of the mixture at room temperature for one hour on a shaker. Bound and free [^3^H]DAMGO were separated by rapid filtration with GF/B filters pre-soaked with polyethyleneimine (1 h at room temperature) under vacuum with a Brandel Cell Harvester followed by washing four times with ice-cold washing buffer (50 mM Tris, pH7.4). Radioactivity on the filters were determined by liquid scintillation counting with a scintillation counter (Perkin-Elmer). The results were presented as specific binding in DPMs/100 μg protein contents.

[^35^S]GTPγS binding was conducted with sonicated tissue preparations (20 μg of proteins), [^35^S]GTPγS (80 pM, 190,000-220,000 dpm) and 80 μM GDP with or without 5 μM DAMGO in a total volume of [^35^S]GTPγS binding buffer (50 mM Tris, 100 mM NaCl, 0.1% BSA, 5 mM MgCl2, and 1 mM EDTA, pH 7.4). Nonspecific binding was defined by incubation in the presence of 10 μM GTPγS. Mixtures were incubated for 60 min at 30°C. Bound and free [^35^S]GTPγS were separated by filtration with GF/B filters under reduced pressure. Radioactivity on filters was determined by liquid scintillation counting.

#### Statistical Analyses

All data were analyzed using GraphPad Prism version 8.2.1 (GraphPad Software Inc., La Jolla, CA). All drug self-administration data were analyzed using a two-way repeated measures analysis of variance (RM-ANOVA), using “day” as the within-subject factor and “sire” as the between-subject factor. Mann-Whitney *U*-tests were used to analyze all progressive ratio infusion data. We only included rats that were patent in these analyses. For the sucrose self-administration experiments, we used an unpaired Student’s *t*-test. Lastly, for the mu-opioid binding studies, data were analyzed using an unpaired Student’s *t*-test. For all data, significance was defined as *p* < 0.05.

## Results

### Paternal morphine exposure does not disrupt maternal behavior or reproductive hormone levels in male or female progeny

Male Sprague Dawley rats had daily 3-hour access to morphine self-administration on a FR1 schedule of reinforcement for 60 days as described previously [32] (**Figure 1A**). Drug-loading behavior refers to the transition from initial controlled consumption to compulsive drug use, which occurs after chronic drug self-administration [40, 41]. Rats that had access to morphine showed increased loading behavior across the 60-day period and control animals exposed to saline did not show any changes in consummatory or loading behavior during that same period (**Supplemental Figure 1**).

The differential allocation hypothesis posits that dams can alter their maternal care based on their assessment of the fitness of their mating partners [42]. In the current experimental design, dams and sires were paired for less than one week. We examined whether maternal behaviors were altered by breeding to a morphine-exposed sire compared to controls in dams that recently produced pups. There were no differences in maternal care comparing dams paired with morphine-treated sires to those bred to saline control males (**Supplemental Figure 2A**), which is consistent with previous reports assessing maternal behaviors between saline- and cocaine-sired litters [33]. In females and males, anogenital distance (AGD) is defined as the distance in millimeters between the anus and the genitals. AGD represents a reliable marker of changes in reproductive and gonadal hormone levels during development [43, 44]. AGD measured at postnatal days 21 and 28 in male and female progeny was unaffected by paternal morphine history (**Supplemental Figure 2B and C**).

### Morphine-sired male but not female offspring show higher motivation for morphine compared to saline-sired controls

Adult (60+ days) male and female F1 progeny were tested for morphine acquisition and reinforcing efficacy at two doses: 0.25 mg/kg/infusion and 0.75mg/kg/infusion. Rats were first trained on a FR1 schedule of reinforcement for 10 days (3hrs/daily) then evaluated for 1 day under a PR schedule. We found that paternal morphine exposure did not alter the acquisition of morphine self-administration in female rats at the 0.25mg/kg dose (**Figure 1B**: effect of sire: *F*_(1, 39)_ = 0.07573, *p* = 0.7846; effect of day: *F*_(9, 351)_ = 1.240, *p* = 0.2690; interaction: *F*_(9, 351)_ = 1.313, *p* = 0.2284). There was also no difference in infusions between saline- and morphine-sired female offspring under a PR schedule of reinforcement (**Figure 1C**; *U* = 29.50, *p* = 0.8176). To determine whether the observed phenotype in female progeny was influenced by the estrous cycle and ovarian hormone fluctuations, we examined morphine self-administration during the estrus and non-estrus phase for both groups. The phase of the estrous cycle had no impact on drug-taking under a PR reinforcement schedule, which was consistent with previous findings [45–48] (**Supplemental Figure 3**). At the 0.75 mg/kg dose, both saline- and morphine-sired female offspring showed increased morphine intake over the acquisition period (**Figure 1D**; effect of day: *F*_(1.634, 52.28)_ = 8.920, *p* = 0.0010). Paternal morphine history had no impact on morphine intake between both groups (effect of sire: *F*_(1, 32)_ = 0.1556, *p* = 0.6959; interaction: *F*_(9, 288)_ = 0.4053, *p* = 0.9318). Paternal morphine exposure also had no effect on the number of morphine infusions earned under a progressive ratio schedule at this higher dose (**Figure 1E**; *U* = 44, *p* = 0.6674). Taken together, these results suggest that paternal morphine history did not impact the self-administration of morphine in female progeny.

In sharp contrast, male morphine-sired offspring self-administered more morphine than controls at the 0.25mg/kg dose (**Figure 1F**; effect of sire: *F*_(1, 36)_ = 10.02, *p* = 0.0031; effect of day: *F*_(9, 324)_ = 1.243, *p* = 0.2677; interaction: *F*_(9, 324)_ = 1.037, *p* = 0.4101). Under a progressive ratio schedule of reinforcement, morphine infusions were significantly higher for morphine-sired males compared to saline-sired males (**Figure 1G;** effect of sire: *U* = 6.500, *p* = 0.0106). At the 0.75mg/kg dose, morphine intake increased significantly over the 10 day acquisition period, but there was no difference in infusions between saline- and morphine-sired progeny (**Figure 1H;** effect of day: *F*_(2.679, 95.84)_ = 3.011, *p* = 0.0393; effect of sire: *F*_(1, 37)_ = 0.0002633, *p* = 0.9871; interaction: *F*_(9, 322)_ = 0.9439, *p* = 0.4870). Paternal morphine history also had no impact on infusions earned under a progressive ratio schedule of reinforcement (**Figure 1I;** effect of sire: *U* = 17.50, *p* = 0.2379). Overall, these data suggest that paternal morphine consumption alters the self-administration and reinforcing efficacy of morphine in morphine-sired male offspring in a dose-dependent manner.

### Paternal morphine taking does not affect self-administration of cocaine, nicotine, or sucrose in F1 male or female progeny

We used separate drug-naive cohorts of F1 offspring to determine whether the intergenerational effect of paternal morphine exposure generalized to other reinforcers. We tested two psychostimulants, cocaine and nicotine, as well as a natural reinforcer-sucrose. Rats were given access to cocaine for 2 hours daily at doses of either 0.25mg/kg/infusion or 0.5 mg/kg/infusion. After 10 days of FR1 cocaine self-administration, rats were switched to a PR schedule to assess the reinforcing efficacy of and motivation for cocaine. For female progeny, a two-way ANOVA revealed that both groups showed increased cocaine self-administration at the 0.25mg/kg dose over the 10-day period and that paternal morphine exposure had no impact on the number of infusions earned (**Figure 2A**; effect of day: *F*_(1.570, 30.69)_ = 12.63, *p* = 0.0003); effect of sire: *F*_(1, 20)_ = 0.003090, *p* = 0.9562); interaction: (*F*_(9, 176)_ = 0.4180, *p* = 0.9243). Saline- and morphine-sired female rats earned a comparable number of cocaine infusions under a PR schedule at this dose (**Figure 2B**; effect of sire: *U* = 51.50, *p* = 0.5950). Similarly, at the 0.5mg/kg dose, all animals displayed increased cocaine self-administration and there was no effect of paternal morphine exposure on the number of infusions earned over 10 days (**Figure 2C**; effect of day: *F*_(1.423, 19.92)_ = 14.03, *p* = 0.0005; effect of sire: *F*_(1, 14)_ = 0.1358, *p* = 0.7180; interaction: (*F*_(9, 126)_ = 1.035, *p* = 0.4159). Paternal morphine history did not have any impact on cocaine taking under a progressive ratio schedule (**Figure 2D**; *U* = 31.50, *p* = 0.9843).

**Figure 2.**
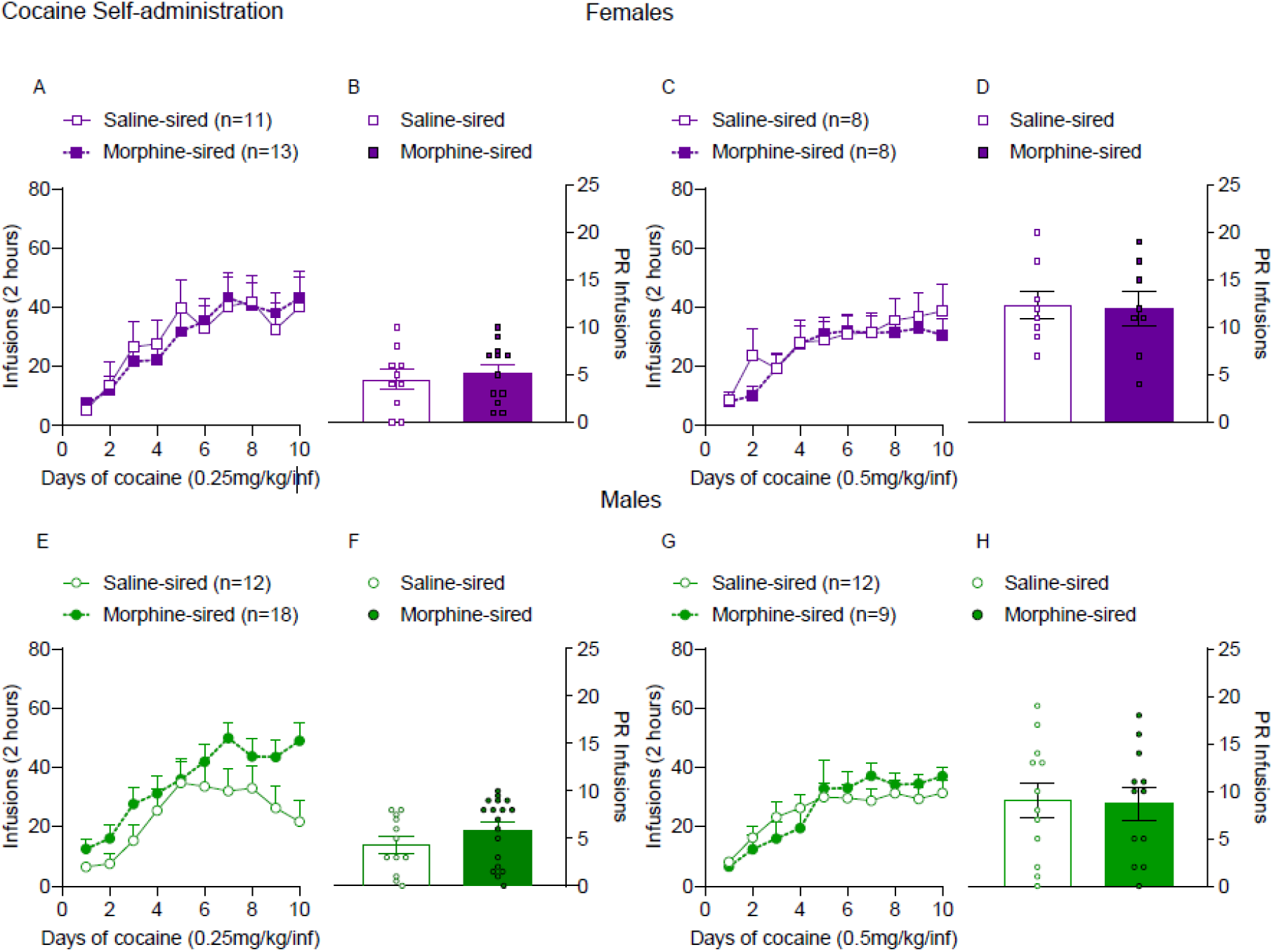
Paternal morphine exposure does not alter cocaine self-administration in female or male progeny. **A.** Paternal morphine consumption did not alter cocaine self-administration in female rats (from 7 saline-exposed sires and 7 morphine-exposed sires) at the 0.25 mg/kg dose. **B.** Saline-sired (from 5 sires) and morphine-sired female rats (from 7 sires) do not show altered cocaine self-administration under a progressive ratio schedule. **C.** Similarly, at the 0.5 mg/kg dose, there was no main effect of paternal morphine exposure on morphine infusions in female rats (from 4 saline-exposed sires and 4 morphine-exposed sires) during the acquisition phase. **D.** Under a progressive ratio schedule, there was no difference in cocaine infusions between saline-sired (from 4 saline-exposed) and morphine-sired female rats (from 4 morphine-exposed sires). **E.** Paternal morphine consumption did not alter cocaine self-administration in male rats (from 7 saline-exposed sires and 9 morphine-exposed sires) at 0.25 mg/kg dose. **F.** Under a progressive ratio schedule of reinforcement, both saline-sired (from 7 saline-exposed sires) and morphine-sired male rats (from 9 morphine-exposed sires) self-administered similar amounts of cocaine at the low dose. **G**. At the higher dose of cocaine (0.5 mg/kg), there was no main effect of paternal morphine on infusions during the acquisition phase in male rats (from 6 saline-exposed sires and 6 morphine-exposed sires). **H.** There was no difference in cocaine infusions between saline-sired (from 6 saline-exposed sires) and morphine-sired male rats (from 6 morphine-exposed sires) under a progressive ratio schedule of reinforcement. Data are expressed as the mean ± S.E.M. *p < 0.05.

Repeated measures ANOVA revealed that the number of cocaine infusions earned increased over the 10-day FR1 acquisition period at the 0.25mg/kg/infusion dose for both groups of F1 males but that paternal morphine history did not affect cocaine self-administration (**Figure 2E**; effect of day: *F*_(2.053, 55.66)_ = 15.45, *p* = 0.0001); effect of sire: *F*_(1, 28)_ = 2.907, *p* = 0.0993); interaction: *F*_(9, 244)_ = 1.133, *p* = 0.3397). Furthermore, under a PR schedule of reinforcement, saline-sired and morphine-sired male offspring earned similar amounts of cocaine (**Figure 2F**; *U* = 73, *p* = 0.1407). At the 0.5 mg/kg/infusion dose, both saline- and morphine-sired males showed an increase in cocaine infusions earned over the course of 10-days, but paternal morphine-history did not have an impact on cocaine consumption (**Figure 2G**; effect of day: *F*_(1.385, 24.94)_ = 22.50, *p* < 0.001; effect of sire: *F*_(1, 18)_ = 0.02578, *p* = 0.8742; interaction: *F*_(9, 162)_ = 1.896, *p* = 0.0559). Paternal morphine history also had no impact on the total cocaine infusions earned under a progressive ratio schedule at this dose (**Figure 2H**; *U* = 70, *p* = 0.9211). Taken together, these data indicate that cocaine self-administration was not impacted by paternal morphine self-administration in in male of female offspring.

For nicotine self-administration, paternal morphine exposure did not alter acquisition (**Supplemental Figure 4A**) or motivation to earn nicotine on a progressive ratio schedule (**Supplemental Figure 4B**) in male progeny. Moreover, self-administration of the natural reward sucrose was not impacted by paternal morphine history in female (**Supplemental Figure 4C**) or male (**Supplemental Figure 4D**) progeny. Consistent with this finding, sucrose preference, a measure of depressive-like behavior was also unaffected by a paternal history of opioid consumption in female (**Supplemental Figure 5A**) and male (**Supplemental Figure 5B**) offspring.

Opioid consumption is thought to be initially driven by the appetitive and reinforcing properties of this class of drug. After chronic consumption, withdrawal-related symptoms and negative reinforcement can also account for the dysregulated and compulsive use that characterizes opioid use disorder [49–52]. We tested precipitated withdrawal in dependent female and male progeny. Animals were injected with morphine (10mg/kg, s.c.) twice per day for 5 days and withdrawal was precipitated by a single dose of naloxone (2mg/kg) (**Supplemental Figure 6A**). Female morphine-sired offspring (**Supplemental Figure 6B**) showed similar withdrawal-like signs compared to controls. Paternal morphine exposure did no impact withdrawal-like signs in male progeny either (**Supplemental Figure 6C**).

### Paternal morphine exposure increases mu-opioid receptor expression in the VTA, but not in the NAc, in male offspring

Brain regions were collected from drug-naïve adult saline- and morphine-sired female and male rats to examine the molecular correlates of the observed increased reinforcing efficacy of morphine in F1 progeny of saline- and morphine-treated sires. Morphine-sired male offspring had significantly greater expression of mu-opioid receptors assessed by [^3^H] DAMGO binding in the VTA compared to male saline-sired controls (**Figure 3A;** *t*_26_=2.653, p = 0.0134). There was no difference in mu-opioid expression in the NAc of drug-naïve male offspring of saline- and morphine-treated sires (**Figure 3B;** *t*_18_=0.2093, p = 0.8366). VTA mu-opioid receptor expression in female offspring was not impacted by paternal morphine history (**Supplemental Figure 7A)**. Signal transduction associated with G-protein activation following MOR activation, assessed with DAMGO-stimulated [^35^S]GTPγS binding, was not changed in the VTA of female or male progeny (**Supplemental Figure 7B&C)**. Taken together, these results demonstrate the paternal morphine exposure alters the expression of the mu-opioid receptor, specifically in the VTA of drug-naïve morphine-sired males.

**Figure 3.**
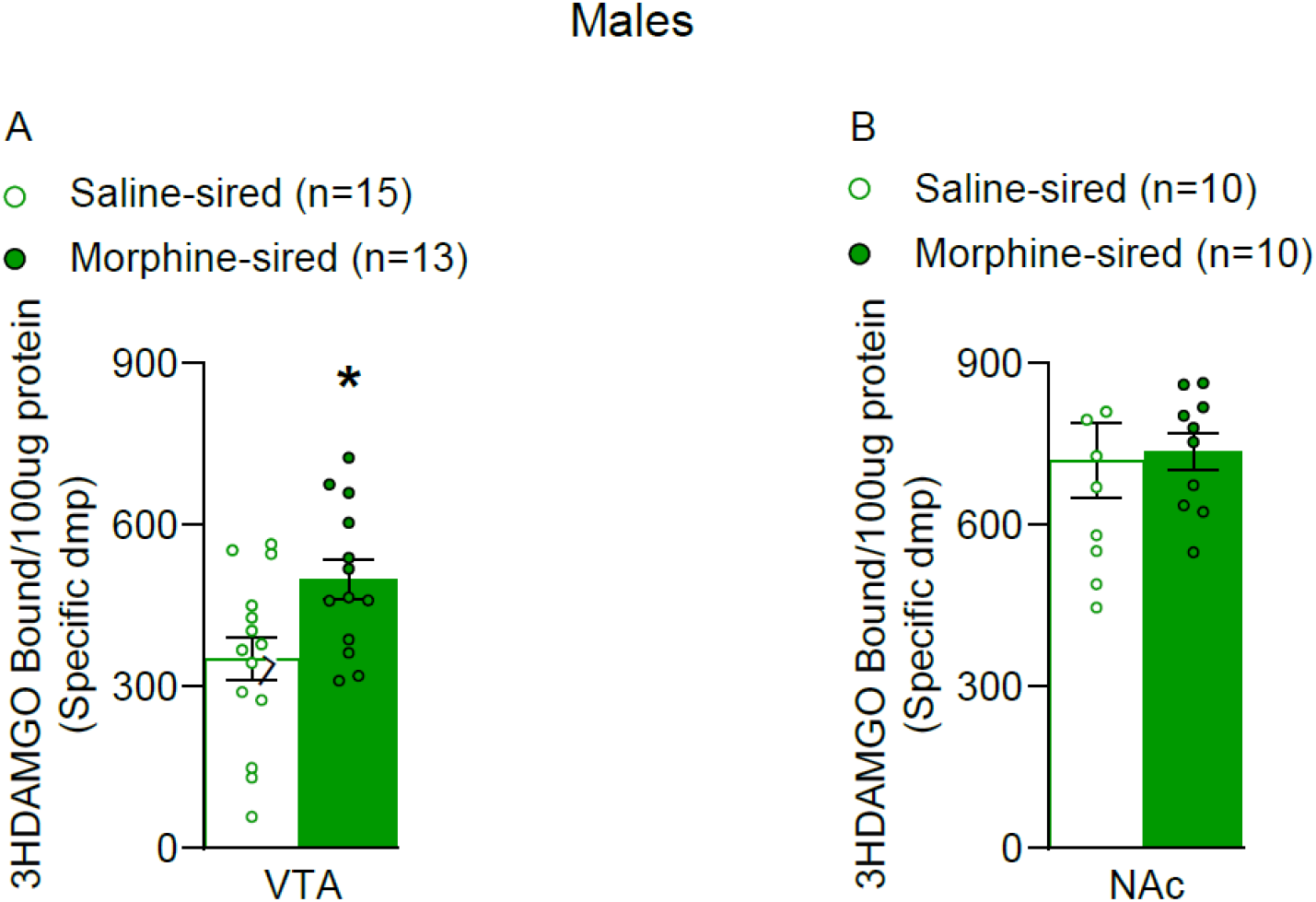
Increased mu-opioid receptor expression in the VTA, but not the NAc of male offspring induced by paternal morphine exposure. Mu-opioid receptor expression in the VTA **(A)** and NAc (**B**)of drug-naïve male rats derived from sires exposed to either saline or morphine. There is significantly greater opioid receptor binding in the VTA of morphine-sired male offspring compared to saline-sired controls. Opioid receptor expression was similar in the NAc of saline-sired and morphine-sired male offspring.

## Discussion

Our results demonstrate that chronic paternal morphine consumption dose dependently increased the reinforcing efficacy of morphine in male but not female F1 progeny. This phenotype did not generalize to other reinforcers, evidenced by the fact that cocaine, nicotine, or sucrose self-administration were all comparable in morphine- and saline-sired progeny. The increased sensitivity to the reinforcing properties of morphine in male offspring produced by morphine-treated sires was associated with increased mu-opioid receptor expression in the VTA. Overall, these results have potential implications for the children of fathers who are chronically exposed to morphine.

### Paternal morphine exposure caused increased morphine-taking in male but not female progeny

Previous multigenerational studies of paternal morphine exposure have yielded conflicting results regarding alterations in reward sensitivity in the next generation [29–31, 53]. Twenty-one days of non-contingent oral morphine treatment in either dams or both parents followed by a 10-day washout period was sufficient to increase morphine preference in progeny on two-bottle choice paradigms. However, paternal exposure alone did not produce this effect. Methamphetamine choice was not impacted in any of the groups. Interestingly, morphine conditioned place preference was blunted when either dams, sires or both were treated with morphine prior to conception. Methamphetamine conditioned place preference was not impacted in any of the groups [53]. When both parents were exposed to morphine for 2 weeks with a month of washout prior to mating, progeny also had increased preference for morphine on a two-bottle choice paradigm, which is consistent with some of the aforementioned and present results. Paternal exposure alone was not sufficient to produce these effects using this regimen of exposure and exercise mitigated the reported phenotype produced by exposure to both parents [30]. In another study, sires received a non-contingent escalating regimen of morphine during adolescence (PND 30-40) and were bred to drug-naïve females as adults. The resulting male and female progeny showed reduced cocaine self-administration and diminished motivation for cocaine. Female first-generation offspring also showed increased morphine intake and increased motivation for oxycodone self-administration. This bidirectional behavioral phenotype was accompanied by increased BDNF expression in the prefrontal cortex of adult progeny. In sharp contrast, adult male offspring produced by sires exposed to morphine during adolescence were resistant to the rewarding effects of low doses of morphine on a conditioned place preference paradigm. This decreased in morphine preference was accompanied by reduced spontaneous burst firing of VTA dopaminergic cells. Here, we chose to focus on the paternal lineage to circumvent some of the potentially confounding factors associated with maternal opioid exposure using a volitional model of morphine self-administration during adulthood. We chose 60 days of exposure to cover the duration of spermatogenesis in rats and sires continued to self-administer morphine to avoid any withdrawal-mediated impact on future generations. Importantly we report no impact on maternal behavior, reproductive hormone levels and have previously shown that litter size is not impacted by this manipulation. Our findings add to this growing body of literature that the regimen, timing and duration of morphine exposure has a profound impact on the resulting phenotypes related to drugs of abuse and reward processing in the next generation of animals.

### Increased mu-opioid receptor expression in the VTA of drug-naive morphine-sired male progeny

Morphine binds to mu-opioid receptors located on GABAergic interneurons in the VTA to ultimately increase dopamine neurotransmission into the nucleus accumbens to mediate reward and motivation [54–59]. We found that drug-naïve morphine-sired male offspring have significantly increased selective agonist DAMGO binding to mu-opioid receptors in the VTA but not in the NAc. This effect is in line with increased morphine self-administration behavior in these animals, suggesting altered sensitivity to morphine in morphine-sired males may be partly explained by increased binding to mu-opioid receptors in the VTA. However, modifications in opioid-receptor binding was not associated with significant changes in mu-opioid receptor activation of the G-protein as measured with agonist-stimulated GTP gammaS binding. In its inactive state, G-protein alpha subunits has a relatively high affinity for GDP over GTP, whereas activation of mu-opioid receptors by morphine shifts the alpha subunit into a higher affinity for GTP versus GDP, leading to changes in protein function or gene transcription [60].Thus, our results could be due to differential activation of signaling pathways that are regulated by opioid receptors [61]. A previous report in Sprague Dawley rats found that morphine exposure in dams resulted in increased levels of mu-opioid receptors in the NAc and decreased levels in the VTA [62]. This further emphasizes the divergent effects of prenatal exposure coming from the paternal versus maternal lineage as well as the importance of the timing of exposure (adolescence vs. adulthood) in the parental generation on molecular and behavioral endpoints in subsequent generations.

### Intergenerational transmission of paternal morphine history

The current work provides evidence for intergenerational transmission that impact reward-related behavior and physiology in offspring. Although the mechanism by which paternal morphine exposure alters reward sensitivity in offspring is not yet understood, other researchers have demonstrated multi- and transgenerational effects resulting from stress [63, 64], diet [5], and toxins [65]. These effects have also been reported in humans [66–68] but is a matter of debate, mainly because of confounding ecological, genetic, and cultural factors. DNA methylation [65, 69, 70], histone post translational modifications [33], and small non-coding RNAs [3, 63, 71] in gametes have been identified as potential carriers of environmental information across generations. Given that mu-, delta-, and kappa-receptors are present in germ cells [72, 73] and the fact that they play an important role in spermatogenesis, it is plausible that chronic morphine exposure in sires caused epigenetic reprogramming events that ultimately influence the developmental trajectory of molecular pathways in offspring brain regions that play a critical role in reward signaling. Future work must be focused on uncovering the molecular mechanisms underlying intergenerational transmission of morphine exposure from father to offspring.

## Conclusions

Here, we showed that paternal morphine exposure impacts reward sensitivity of morphine in male but not female progeny as assessed using self-administration. These findings add to a growing body of literature that suggest the role of epigenetic modifications in reprogramming the male germline and thus impact subsequent generations. Future studies are needed to identify the specific molecules involved in the transmission of paternal morphine experiences. Together, these findings have potential consequences for the children of fathers living with opioid use disorders.

## Acknowledgement

We thank Dr. Ellen Unterwald for input on the manuscript and studies. This work was supported by NIH/NIDA K01 DA039308 (MEW), DP1 DA046537 (MEW), T32 DA007237 (ABT; Unterwald EM, PI), P30 DA013429 (LYLC) and R01 DA041359 (LYLC), R01 DA037897, R21 DA039393 and R21 DA045792. We thank Melissa Knouse and Arthur Thomas for help with these studies.

## Notes

### Competing Interest Statement

The authors have declared no competing interest.

