## Supplemental Information for "Paternal morphine exposure enhances morphine self-administration and induces region-specific neural adaptations in reward-related brain regions of male offspring"

Toussaint et al.

**Supplementary Material and Methods:**

**Vaginal cytological sample collection and analysis**

Given the known variations in drug-taking behavior across the estrous cycle in female rats, a separate group of saline-sired and morphine-sired female rats were used to evaluate whether differences in the estrous cycle may influence morphine self-administration. Samples were collected using cotton tips (Medline Industries Inc. Northfield, IL) moistened with saline. One cotton tip per rat was inserted into the vaginal orifice (<1 cm), gently rotated 360 degrees, and removed. The cotton tip was swabbed onto a microscope slide that corresponded with the day of collection. In order to cover three full estrous cycles, we collected vaginal cytology data prior to the start of each self-administration session for 4 days total. After collection, slides were analyzed under a microscope using 40x magnification. Specific cell types that uniquely correlate to each phase of the estrous cycle were identified, and data was used to determine which phase of the cycle each animal was in for every day of the experiment. Following cytological analysis of estrous cycle data, the number of days for each cycle of F1 female offspring was counted to determine the average cycle length of both groups. The length of a singular cycle was determined by designating proestrus as day 1, and then counting the number of days until the next occurrence of proestrus but excluding that day from the count [1]. Female rats self-administered morphine at 0.25mg/kg dose under the following schedule: 10 days of FR1, 2 days of FR3, and 4 days of PR. Each session lastedlasted 3-hrs, 3-hrs, and 6-hrs, respectively.

**F1 Nicotine Self-Administration**

Nicotine self-administration experiments were performed as described previously [2-7]. Following a 7-day recovery from intravenous catheter implantation, a separate drug naïve cohort of F1 offspring self-administered intravenous nicotine (0.03 mg/kg nicotine/59 µl saline, infused over 5 sec) on an FR1 schedule of reinforcement for 10 days, then were switched to a PR schedule. Each nicotine infusion was paired with a light cue and followed by a 20-second timeout period, during which drug infusions were not available but lever presses were recorded. For these experiments, nicotine hydrogen tartrate salt (Sigma) was dissolved in sterile 0.9% saline (pH adjusted to 7.4 ± 0.5 with sodium hydroxide).

**F1 Sucrose Self-Administration**

A separate group of drug-naïve adult littermates were allowed to lever press for sucrose pellets (TestDiet) on a FR1 schedule of reinforcement for 10 consecutive days during daily 1-hr sessions. The main purpose of this experiment was to test for any deficits in operant learning in offspring. Each session lasted 1-hr with no limit on the number of sucrose pellets earned.

**Two Bottle Choice Sucrose Self-Administration**

Animals were singly housed, and their cages were fitted with two bottles (50 mL Falcon tubes, VWR international) containing tap water for at least two days of habituation. Food was removed prior to the start of habitation and testing. Body weight and mass of the water were recorded before each habituation session began, then again 6-hrs later when it ended. At the end of habituation day 2, if animals showed a preference greater than 75% to a particular bottle location (Preference = [sucrose consumption left or right side]/[sucrose consumption left side + sucrose consumption right side] * 100), then sucrose solution was assigned to the opposite side. On the test day, each rat had a choice between two bottles: one filled with tap water and the other with sucrose dissolved in water. Sucrose solutions were 0.25% and 0.5%, respectively. Bottles were positioned equidistant from one another and were randomly alternated to avoid side bias. At the end of testing, the volume of sucrose and water consumption was measured for each rat, and percent sucrose preference was calculated as follows: sucrose preference (%) = (sucrose consumption)/(sucrose consumption+ water consumption) * 100 [8].

**Precipitated Morphine Withdrawal Induced by Naloxone**

Female and male F1 rats were handled for 2 min each day for 5 days leading up to testing. In order to induce morphine dependence, researchers administered morphine injections to drug-naïve saline- and morphine-sired progeny using the following regimen: morphine (s.c.) at 10 mg/kg twice daily for 5 consecutive days. Rats remained in their home cage throughout the experiment. Twenty-four hours after the last morphine injection, rats were given naloxone (2 mg/kg, s.c.) and withdrawal signs were recorded for 30 min according to a modified version of the Gellert & Holtzman scale [9]. Withdrawal-like behaviors including escape attempts, wet-dog shakes, abdominal constrictions, face wiping, grooming, abnormal posture, teeth chattering, and paw shakes. These were counted as the number of events occurring during the total test time. The overall severity of withdrawal from morphine was calculated by totaling the number of times rats exhibited a withdrawal sign.

### Statistical Methods

All data were analyzed using GraphPad Prism version 8.2.1 (GraphPad Software Inc., La Jolla, CA). For sire loading behavior, we used a RM-ANOVA, using “time” as the within-subject factor and “treatment” as the between subject factors. Sidak’s post-hochoc test was used to determine significant difference in drug-loading behaviors over time. For maternal behavior, we used a Chi-squared statistic. For the anogenital distance measures, we used a RM-ANOVA, using “postnatal day” as the within-subject factor and “sire” as the between-subject factor. The two-bottle sucrose preference test was analyzed using a repeated measures two-way ANOVA, with “postnatal day” as the within-subject factor and “sire” as the between-subject factor. For the precipitated withdrawal study, we used a RM-ANOVA, using “withdrawal-like behaviors” as the within-subject factor and “sire” as the between-subject factor. For all data, significance was defined as $p < 0.05$.

**Supplementary Results:**

**Drug-loading behavior in morphine-exposed sires**

We recorded the number of morphine (or saline) infusions sires earned during the first 20 minutes of each session starting on day 1 and every 10th day until day 60 to determine drug-loading behavior. Two-way ANOVA (within-subject factor = self-administration session; between-subjects factor: sire treatment) revealed that rats pressing for morphine, but not those exposed to saline increased the number of infusions earned during the first 20 minutes across the 60 day session (**Supplemental Figure 1**: effect of self-administration session: $F_{(5.085,\ 504.2)}$= 9.671, p<0.0001, effect of sire treatment: $F_{(1,\ 106)}$=8.046, p<0.0055, interaction: $F_{(6.595)}$=4.120, p<0.0005). Sidak's post hoc tests demonstrated a significant increase in the number of infusions morphine-exposed sires took across the 60-day session compared to day 1, while saline-exposed rats took similar amounts of infusions across each session. These results demonstrate that rats with a history of morphine were faster to escalate their consumption to a level that is very stable, suggesting a new set point that regulates morphine self-administration and is a key feature in the motivation for taking opiate drugs.

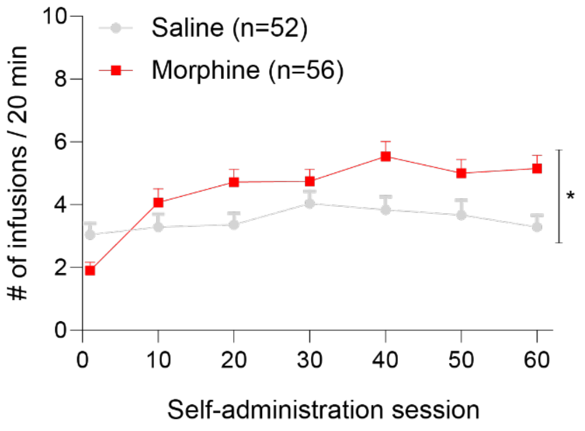


**Supplemental Figure 1**. **Drug-loading behavior is increased in rats chronically exposed to morphine**. Infusions for saline-exposed and morphine-exposed sires taken within the first 20 minutes starting on day 1 and every 10$^{th}$ day until day 60 were quantified and analyzed. Only morphine-exposed rats displayed a significant difference in drug-loading behaviors over time compared to day 1. Data are expressed as the mean ± S.E.M. *$p < 0.05$.

### Maternal care and anogenital distance are unaffected by paternal morphine exposure

The differential allocation hypothesis posits that changes in maternal-care investment directed to offspring may arise through differential breeding with undesirable sires [10-12]. To determine whether paternal morphine history impacted maternal care directed towards F1 pups, we made blinded observations twice daily for 1hr from PND3 to 14. Maternal care behaviors included: dam licking and grooming any pup; dam nursing in an arched position, blanket or passive posture; and dam off pups. The percentage of time dams spent engaged in these different maternal behaviors was calculated. Maternal care behavior did not differ for dams bred to either saline-exposed or morphine-exposed sires (**Supplemental Figure 2A**: effect of breeding: $\chi^2(4) = 0.2874$, p = 0.9906).

Next, anogenital distance (AGD), which is a sexually dimorphic landmark and a sensitive biomarker used to assess reproductive hormone abnormalities in animals, was measured to determine the potential impact of paternal morphine exposure on anogenital development in offspring. Measurements were taken at PND21 and 28 in female and male offspring. AGDs were significantly shorter in females than in males on both PND21 and 28, regardless of their siring condition (**Supplemental Figure 2B and C:** female and male saline-sired offspring: effect of postnatal day: $F_{(1, 78)} = 418.1$, $p < 0.0001$; effect of sex: $F_{(1, 78)} = 631.2$, $p < 0.0001$; interaction: $F_{(1, 78)} = 32.22$, $p < 0.0001$; female and male morphine-sired offspring: effect of postnatal day: $F_{(1, 93)} = 582.8$, $p < 0.0001$; effect of sex: $F_{(1, 93)} = 705.7$, $p < 0.0001$; interaction: $F_{(1, 93)} = 37.50$, $p < 0.0001$). Furthermore, the AGD was longer at PND28 compared to PND21 in females, and siring had no impact on the anogenital distance at either developmental time point (**Supplemental Figure 2B:** effect of postnatal day: $F_{(1, 70)} = 556.1$, $p < 0.0001$; effect of sire: $F_{(1, 70)} = 3.363$, $p = 0.0709$; interaction: $F_{(1, 70)} = 1.131$, $p = 0.2913$). For male offspring, the anogenital distance was longer at PND28 compared to PND21 and siring had no effect on the anogenital distance at either developmental time point (**Supplemental Figure 2C:** effect of post-natal day: $F_{(1, 101)} = 681.4$, $p < 0.0001$; effect of sire: $F_{(1, 101)} = 1.200$, $p = 0.2760$; interaction: $F_{(1, 101)} = 0.2557$, $p = 0.6142$). Taken together, these results suggest that paternal morphine exposure does not impact maternal care directed towards pups, nor does it alter hormonal development during adolescence in female or male progeny.

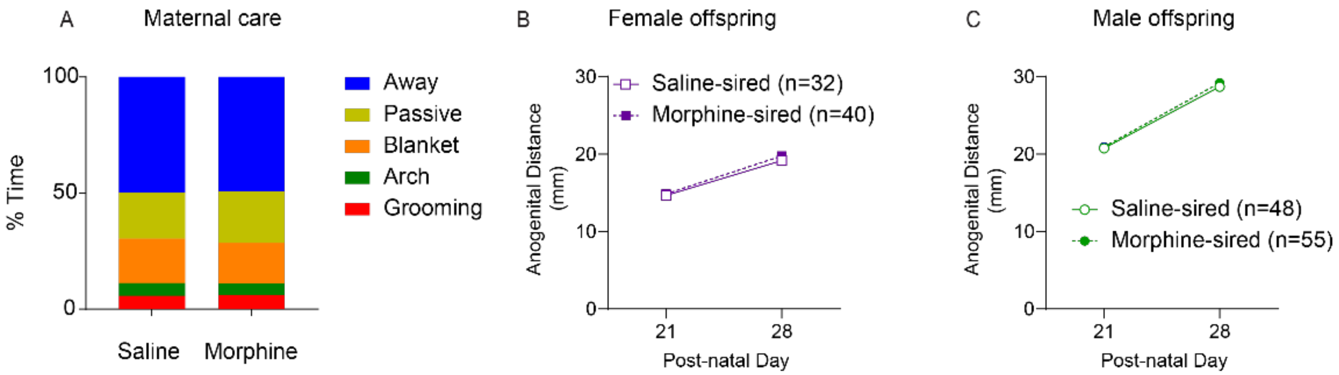


**Supplemental Figure 2. Maternal care directed towards pups and anogenital distance are unaffected by paternal morphine exposure. A.** Data were collected from saline-paired (n=9) and morphine-paired (n=6) dams. Researchers were blind to the sire breeding condition. Maternal care behavior did not differ for dams bred to either saline-exposed or morphine-exposed sires. **B, C.** Anogenital distance (AGD) measurements collected at PND 21 and 28 from offspring derived from saline-exposed (from 8 sires) or morphine-exposed sires (from 9 sires). AGD at both time points were significantly shorter in female compared to male offspring. Siring had no effect on AGD at either developmental timepoint in female and male progeny. Data are expressed as the mean ± S.E.M. *p < 0.05.

### Estrous cycle does not impact morphine taking under progressive ratio schedule of reinforcement

Given the known variations in drug-taking behavior across the estrous cycle in female rats [13], a separate group of saline-sired and morphine-sired female rats were used to evaluate whether differences in the estrous cycle may influence morphine self-administration at the 0.25 mg/kg dose. Vaginal cytology slides were collected prior to the start of each self-administration session. Breakpoints on a progressive ratio reinforcement schedule were not affected by paternal morphine history or estrous cycle in female progeny (**Supplemental Figure 3** effect of sire: $F_{(1, 17)} = 1.097$, $p = 0.3096$; effect of estrous: $F_{(1, 16)} = 0.002342$, $p = 0.9620$; interaction: $F_{(1, 16)} = 0.4323$, $p = 0.5202$; Saline-sired non-estrus and estrus: $U = 22$, $p = 0.5153$; Morphine-sired non-estrus and estrus: $U = 50$, $p = 0.5075$).

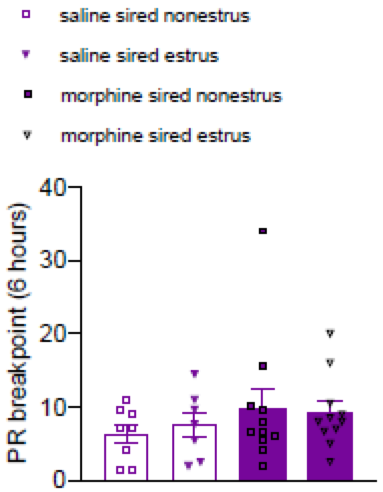


**Supplemental Figure 3. The estrous cycle had no impact on morphine taking under a PR reinforcement schedule.** Vaginal cytology slides were collected prior to the start of each self-administration session for 4 four consecutive days. Analyses were done based on siring and phase of the cycle (nonestrus or estrus). Breakpoints on a PR reinforcement schedule were not affected by paternal morphine history or estrous cycle in female progeny Data are expressed as the mean ± S.E.M. *p < 0.05.

### Paternal morphine exposure does not alter nicotine or sucrose self-administration in F1 offspring

Naïve littermates were used for these experiments. Male saline-sired and morphine-sired offspring had access to nicotine (0.03 mg/kg) via intravenous self-administration on an FR1 schedule for 10 consecutive days. Both groups gradually took higher infusions of nicotine but there was no effect of paternal morphine on self-administration (**Supplemental Figure 3A:** effect of day: $F_{(1.575, 22.04)} = 11.29$, $p < 0.001$; effect of sire: $F_{(1, 14)} = 0.5201$, $p = 0.4827$; interaction: $F_{(9, 126)} = 0.9816$, $p = 0.4586$). Under a progressive ratio schedule of reinforcement, saline-sired and morphine-sired male rats self-administered similar amounts of nicotine infusions (**Supplemental Figure 4B:** effect of sire: $U = 165.5$, $p = 0.3518$), suggesting no impact on the reinforcing efficacy of nicotine.

For sucrose self-administration, both groups of female progeny self-administered an increasing number of pellets over time but there was no main effect of paternal morphine exposure (**Supplemental Figure 5C:** effect of day: $F_{(3.342, 46.79)} = 42.82$, $p = 0.0001$; effect of sire: $F_{(1, 14)} = 2.275$, $p = 0.1537$; interaction: $F_{(8, 112)} = 1.070$, $p = 0.3891$). Similarly in male offspring, both groups took increasingly more sucrose pellets over time but there was no main effect of paternal morphine exposure (**Supplemental Figure 6D:** effect of day: $F_{(2.980, 65.56)} = 11.36$, $p = 0.0001$; effect of sire: $F_{(1, 22)} = 0.7019$, $p = 0.4112$; interaction: $F_{(9, 198)} = 0.4824$, $p = 0.8854$). Taken together, these results demonstrate that paternal morphine history does not impact nicotine self-administration, nor does it produce deficits in operant learning in progeny.

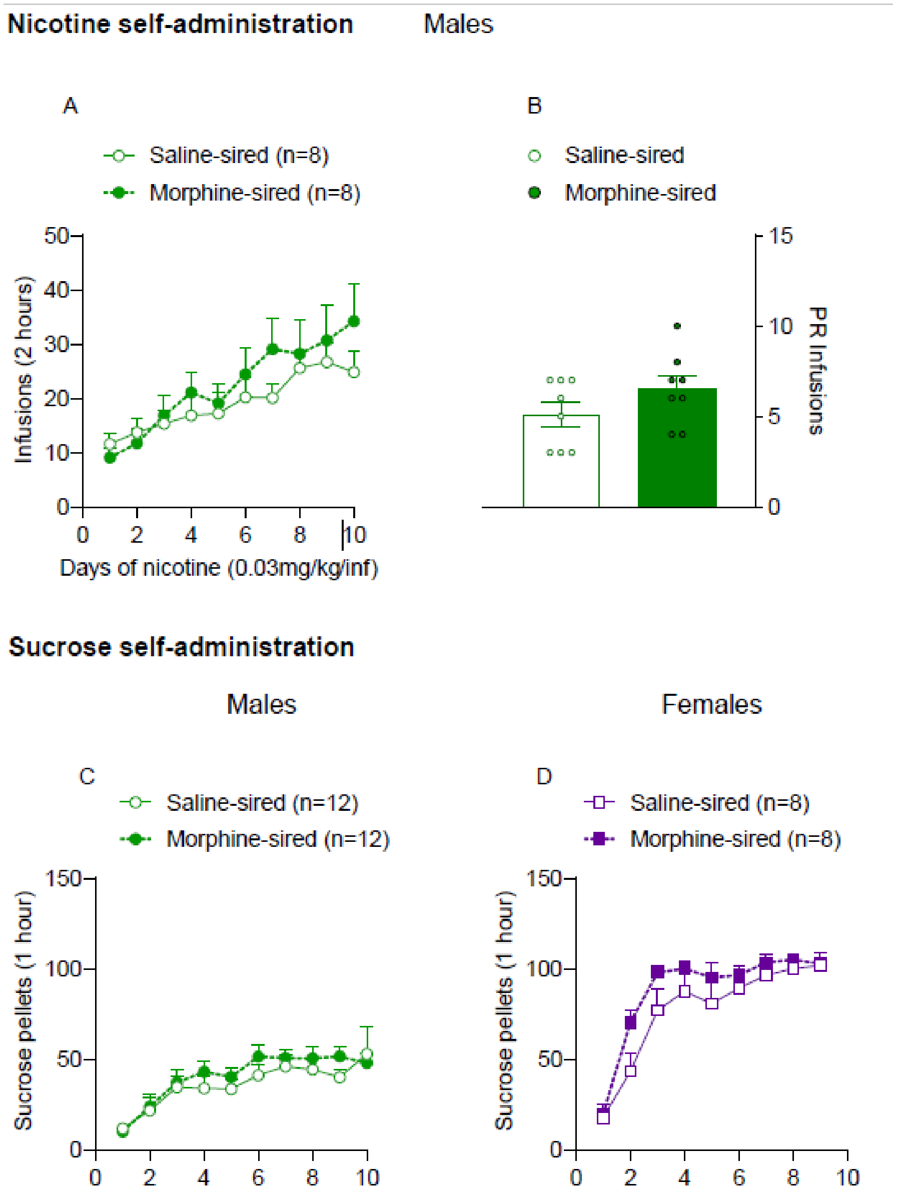


**Supplemental Figure 7.** Paternal morphine exposure does not alter nicotine or sucrose self-administration in F1 offspring. **A.** There was no main effect of paternal morphine exposure on nicotine self-administration in male rats (from 5 saline-exposed sires and 6 morphine-exposed sires). **B.** When switched to a progressive ratio schedule of reinforcement, saline-sired (from 5 saline-exposed sires) and morphine-sired (from 6 saline-exposed sires) male rats self-administered similar amounts of nicotine infusions. **C**. In female progeny (from 4 saline-exposed sires and 4 morphine-exposed sires), there was no main effect of paternal morphine exposure on sucrose taking, which demonstrates that there were no deficits in operant learning. **D.** In male offspring (from 6 saline-exposed sires and 6 morphine-exposed sires), there was no main effect of paternal morphine exposure on deficits in operant learning. Data are expressed as the mean ± S.E.M. *$p < 0.05$.

**Paternal morphine history does not elicit depressive-like behaviors in female and male progeny**

Recent clinical evidence suggests that parental drug use could be a strong risk factor for mental health problems such as depression in children [14]. To investigate whether paternal morphine exposure induced depressive-like behaviors in first-generation offspring, we used a two-bottle sucrose preference test [15]. Paternal morphine exposure had no impact on sucrose preference in female (**Supplemental Figure 8A***:* effect of sire: $t_{21}$ =0.9486, *p* = 0.3536) and male progeny (**Supplemental Figure 9B***:* effect of sire: $t_{24}$=1.023, *p* = 0.3166). Together, these results indicate that paternal morphine history does not produce anhedonic behavior in progeny.

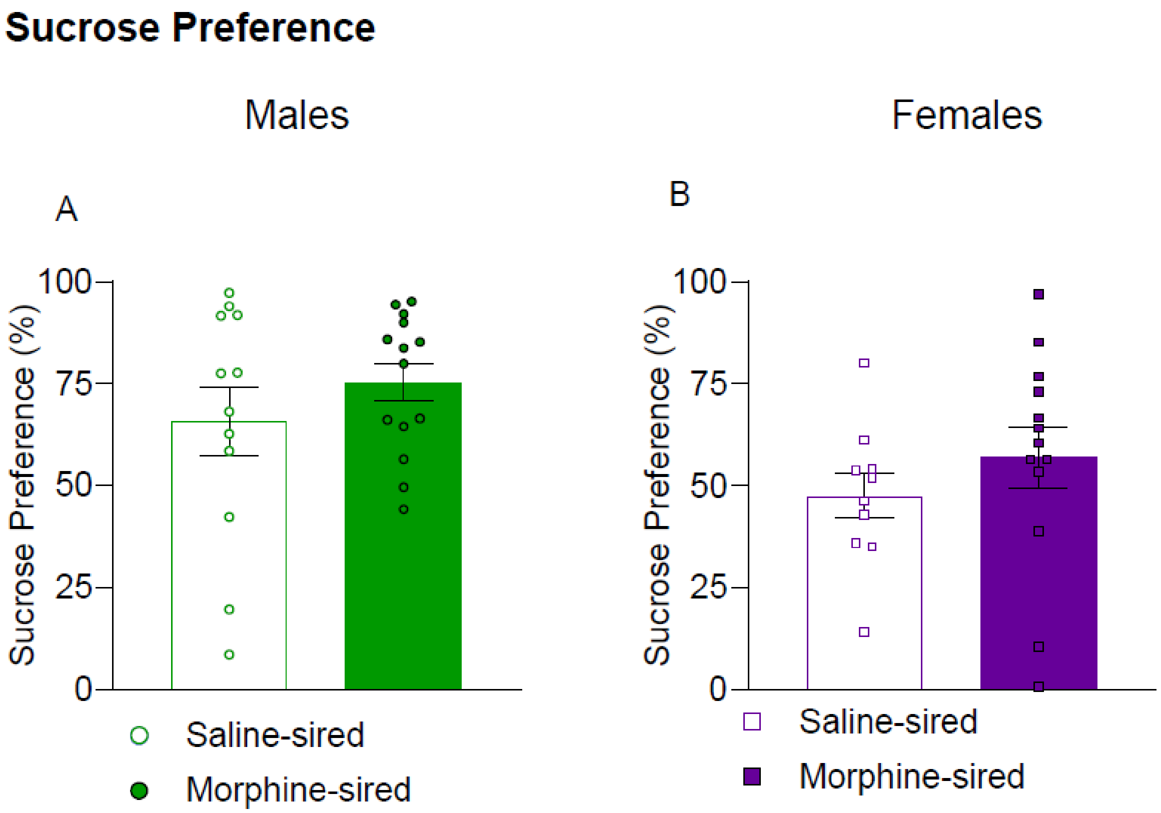


**Supplemental Figure 10**. Paternal morphine history does not elicit depressive-like behaviors in female and male F1 offspring. **A.** Paternal morphine exposure had no impact on sucrose preference in female progeny (from 5 saline-exposed sires and 7 morphine-exposed sires). **B.** Paternal morphine exposure had no impact on sucrose preference in male progeny (from 6 saline-exposed sires and 7 morphine-exposed sires progeny. Data are expressed as the mean ± S.E.M.

**A history of paternal morphine exposure does not impact offspring naloxone-induced withdrawal from morphine**

A separate cohort of rats received non-contingent subcutaneous morphine injections (10mg/kg, s.c., twice daily) for 5 consecutive days. Twenty-four hours after the last morphine injection, rats received naloxone injections (2mg/kg, i.p.) and withdrawal signs were recorded for 30 min (**Supplemental Figure 11A**). For female saline-sired and morphine-sired progeny, paternal morphine consumption did not affect withdrawal-like behaviors (**Supplemental Figure 12B:** effect of withdrawal-like behavior: $F_{(3.491, 34.91)} = 0.8416$, $p = 0.4952$; effect of sire: $F_{(1, 10)} = 0.1025$, $p = 0.7554$; interaction: $F_{(7, 70)} = 0.9434$, $p = 0.4792$).). Similarly for males, withdrawal-like behaviors were not impacted by paternal morphine history (**Supplemental Figure 13C:** effect of withdrawal-like behavior: $F_{(2.033, 20.33)} = 0.8194$, $p = 0.4565$; effect of sire: $F_{(1, 10)} = 0.1889$, $p = 0.6730$; interaction: $F_{(7, 70)} = 0.3642$, $p = 0.9200$). These results demonstrate that paternal morphine history does not alter naloxone-induced precipitated withdrawal in morphine-dependent progeny.

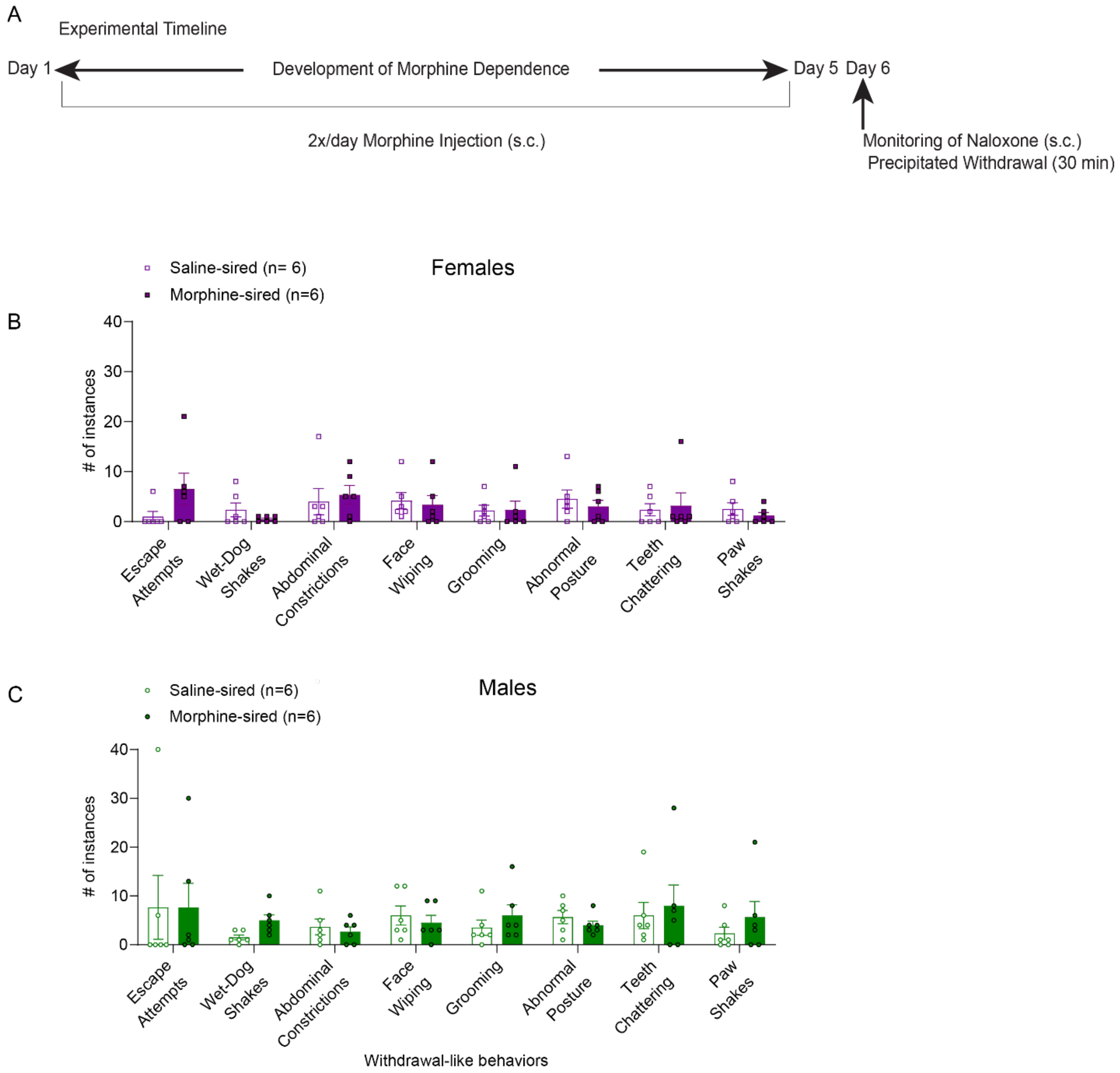


**Supplemental Figure 14**. **A history of paternal morphine exposure does not impact naloxone-induced withdrawal from morphine in offspring**. **A.** Experimental timeline to induce morphine dependence. Adult F1 offspring received morphine (10 mg/kg, s.c., twice daily) for 5 consecutive days. On day 6, offspring were injected with naloxone (2 mg/kg, s.c.) and their withdrawal-like behaviors were monitored for 30 minutes. **B, C.** Paternal morphine consumption did not affect withdrawal-like behaviors in both either saline-sired and or morphine-sired female (from 2 saline-exposed sires and 4 morphine-exposed sires) and or male progeny (from 3 saline-exposed sires and 3 morphine-exposed sires). Data are expressed as the mean ± S.E.M. *$p < 0.05$.

**Paternal morphine history did not impact mu-opioid receptor expression in female progeny, or the MOR agonist DAMGO-stimulated [$^{35}$S]GTPγS binding in male and female progeny**

Adult drug-naïve tissue samples were collected for these analyses. There was no difference between saline-sired and morphine-sired female progeny in mu-opioid receptor expression within the VTA measured with [$^{3}$H]DAMGO binding (**Supplemental Figure 15*A:*** effect of sire: $t_5$=1.205, *p* = 0.2821). MOR-mediated activation of Gα proteins by the selective MOR agonist DAMGO as determined with [$^{35}$S]GTPγS binding in female progeny was also unchanged following paternal morphine history (**Supplemental Figure 16B:** effect of sire: $t_{12}$=0.5457, *p* = 0.5953). Similarly, saline-sired and morphine-sired males show no significant difference in mu-opioid receptor-mediated [$^{35}$S]GTPγS binding within the VTA (**Supplemental Figure 17C*:*** effect of sire: $t_{14}$=1.752, *p* = 0.1017). Taken together, these results demonstrate there is no change in mu-opioid receptor binding and activation within the VTA in female progeny. For males, while morphine-sired offspring had higher levels of the mu-opioid receptor at baseline (see main results), the functional consequence of receptor occupancy, as assessed using [$^{35}$S]GTPγS binding, was not impacted by paternal morphine history.

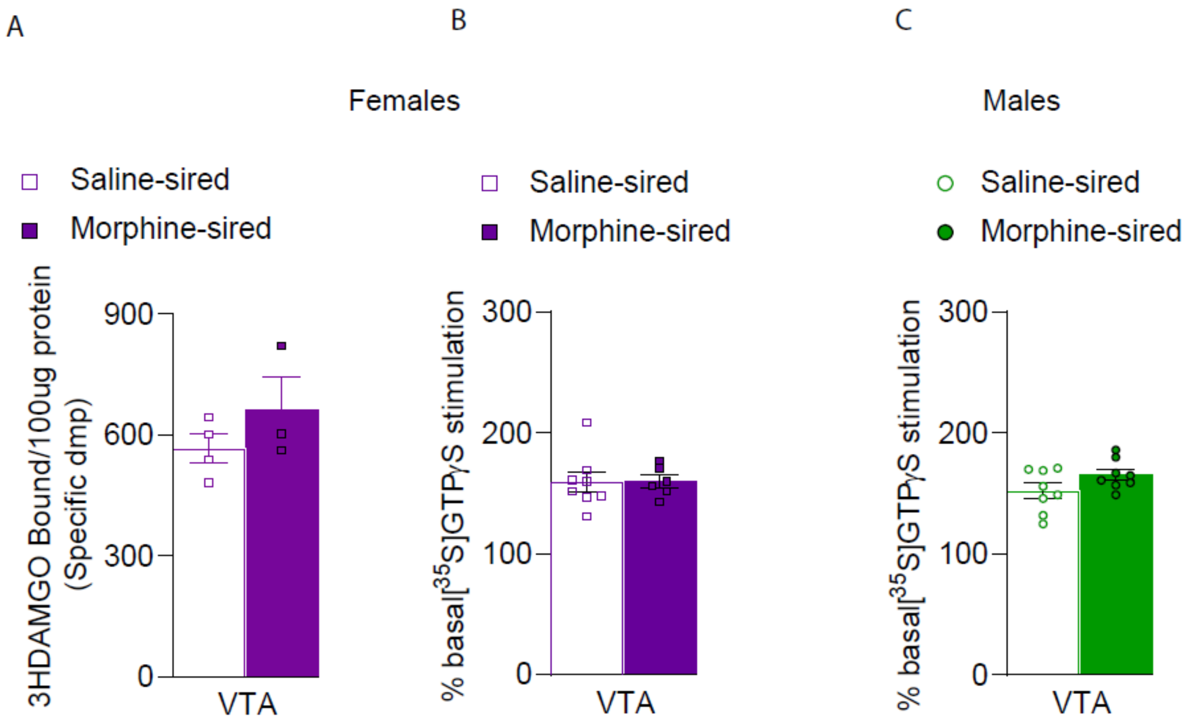


**Supplemental Figure 18**. **Paternal morphine history did not impact mu opioid receptor expression and GTP gamma S activation in female progeny, and GTP gamma S activation in male progeny**. **A.** Drug-naïve saline-sired and morphine-sired female progeny show no significant difference in mu-opioid receptor expression within the VTA. **B**. Saline-sired and morphine-sired female rats show no significant difference in GTP gamma S signal transduction in the VTA. **C**. Saline-sired and morphine-sired male progeny show no significant difference in GTP gamma S signal transduction in VTA. Data are expressed as the mean ± S.E.M. *$p < 0.05$.
